## Supplementary figures and images for "Protein structure and sequence re-analysis of 2019-nCoV genome does not indicate snakes as its intermediate host or the unique similarity between its spike protein insertions and HIV-1"

### Figure 1

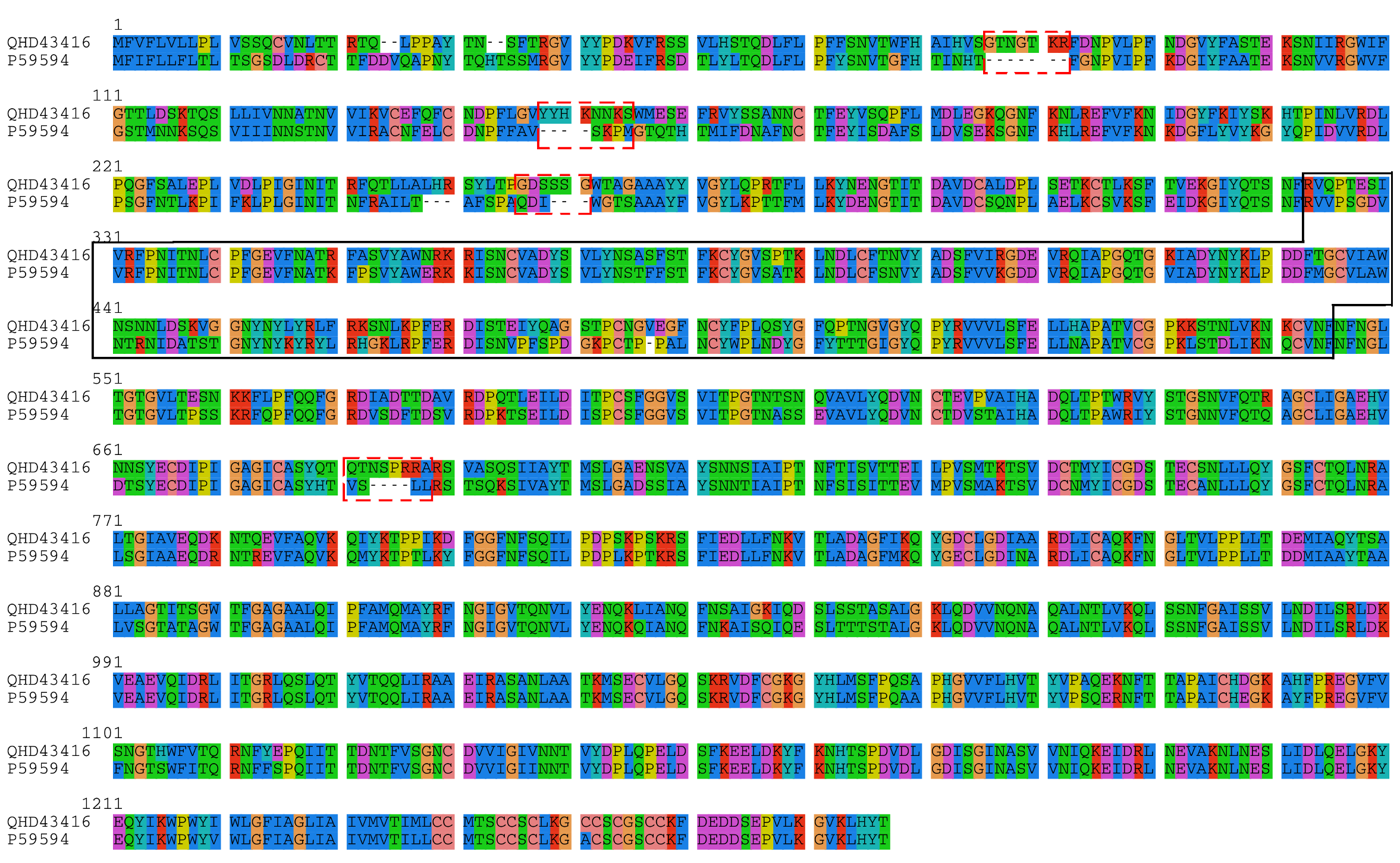

### Figure 2

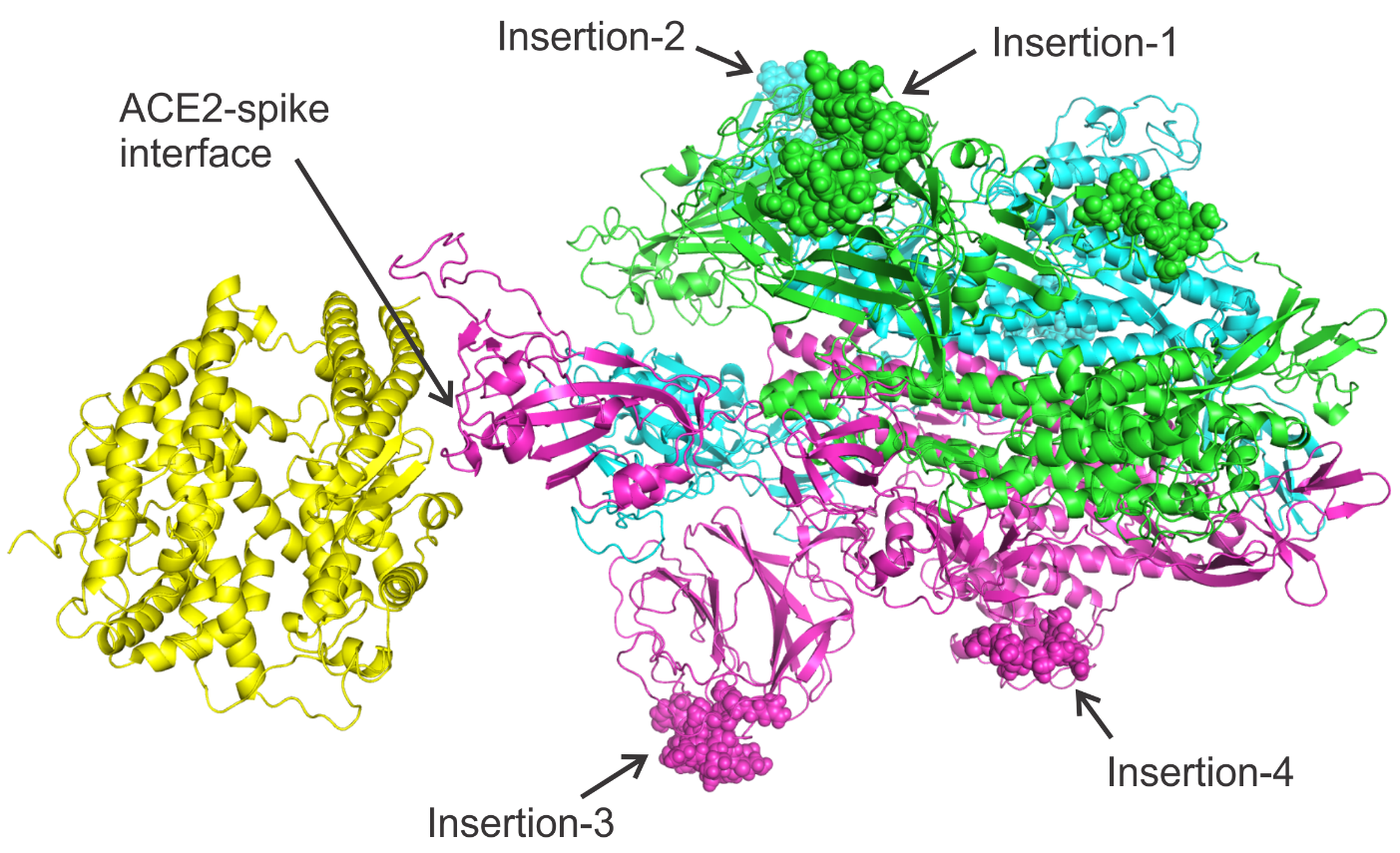

### Figure 3

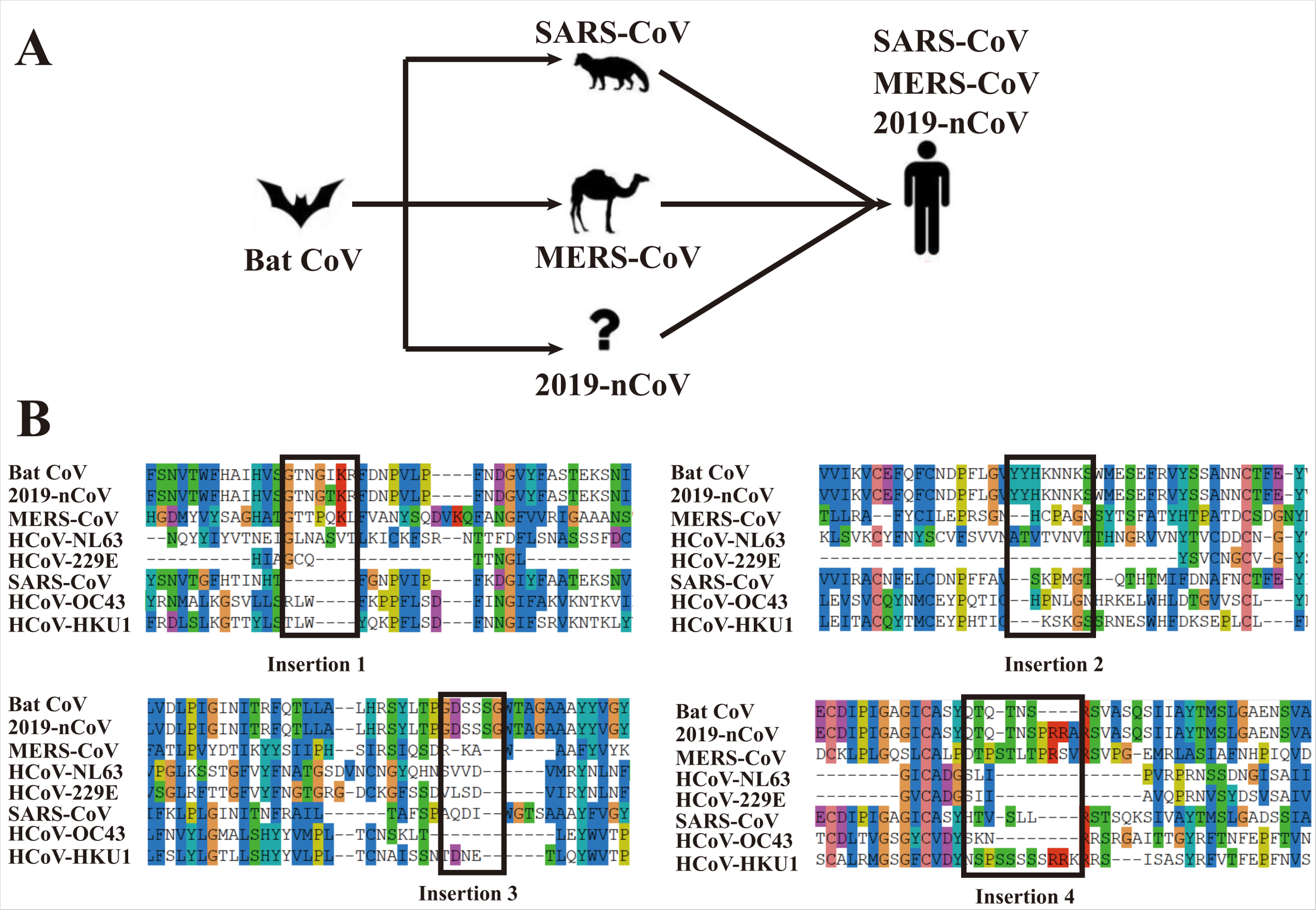

### Figure 4

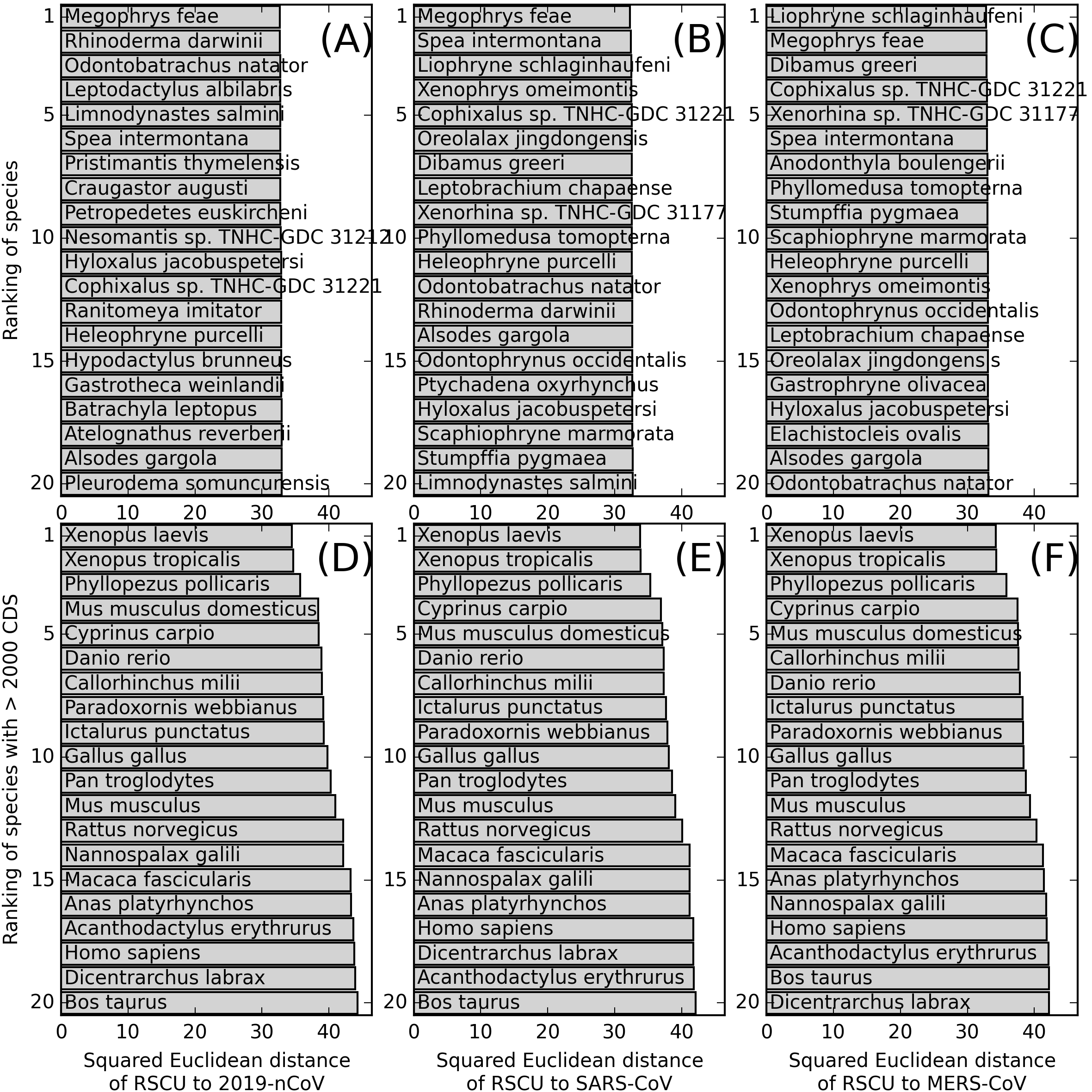
